# Repetitive mild traumatic brain injury causes synergistic effects on mortality

**DOI:** 10.1101/2020.12.22.424055

**Authors:** Ashley M Willes, Tori R Krcmarik, Alexander E Daughtry, Douglas J Brusich

## Abstract

Repetitive mild TBI (rmTBI) events are common in the U.S. However, rmTBI is challenging to study and this contributes to a poor understanding of mechanistic bases for disease following these injuries. We used fruit flies *(D. melanogaster)* and a modified version of the high-impact trauma (HIT) method of TBI to assess the pattern of mortality observed after rmTBI. We found that the pattern of mortality was synergistic after a critical number of injuries, similar to that observed previously at more moderate levels of TBI severity. The identity of cellular and molecular factors which contribute to the synergistic effect on mortality remain unknown, but this model offers a platform for investigation into such factors.

## Description

Millions of Americans suffer traumatic brain injury (TBI) each year (Taylor et al. 2017). The vast majority (~90%) of TBI events in the U.S. are mild traumatic brain injuries, which includes concussions (Cassidy et al. 2004). Mild TBI commonly causes temporary symptoms, but repetitive mild TBI (rmTBI) is associated with long-term consequences which may take years to manifest (Bailes et al. 2013). Current animal models of rmTBI have drawbacks which contribute to the scarcity of evidence for mechanistic bases of dysfunction and disease.

We previously reported an extended fly *(D. melanogaster)* model of TBI based on the high-impact trauma (HIT) method (Katzenberger et al. 2013; Putnam et al. 2019). We showed that repetitive injuries at the relatively moderate deflections of 70° and 80°, but not severe injuries at 90° nor mild injuries at 60°, resulted in a synergistic effect on mortality (Putnam et al. 2019). One potentially confounding issue preventing identification of synergistic effects at 60° was the low mortality rate after the 1-4 injuries administered (Putnam et al. 2019). We chose to further investigate the relationship between mortality and rmTBI using the 60° deflection by extending the injury number to 5-12 HITs.

The main outcome measured using the HIT model of TBI in flies is the mortality index at 24 hours (MI24) (Katzenberger et al. 2013). The pattern of mortality across varied HIT numbers can be assessed by dividing each MI24 value by its respective HIT number (MI24/HIT) (Katzenberger et al. 2013; Putnam et al. 2019). If mortality from each HIT is additive then the MI24/HIT values should be equivalent, but, if mortality is synergistic, then the MI24/HIT values should become increasingly large with HIT number and differ. Neither, *w^1118^* nor *y^1^w^1^* flies showed any differences in MI24/HIT values across 5-8 HITs (Fig. 1A, 1B respectively). However, by 10 HITs both genotypes showed larger MI24/HIT values than for 5 or 6 HITs, and we found a more pronounced effect at 12 HITs (Fig. 1A, 1B). To more fully assess the pattern of mortality across injury number we checked the MI24/HIT data using a line fit. When we used data across only 5-8 HITs neither *w^1118^* nor *y^1^w^1^* flies deviated from fit to a zero-slope line *(w^1118^:*p = 0.34; *y^1^w^1^:* p = 0.45), consistent with additive effects as seen previously. However, inclusion of data from 5-12 HITs for each of *w^1118^* and *y^1^w^1^* resulted in positive slope lines that significantly deviated from zero-slope lines (Fig. 1C, 1D respectively; *w^1118^:* p = 0.004; *y^1^w^1^:* p = 0.006), consistent with a synergistic relationship.

**Figure 1:**
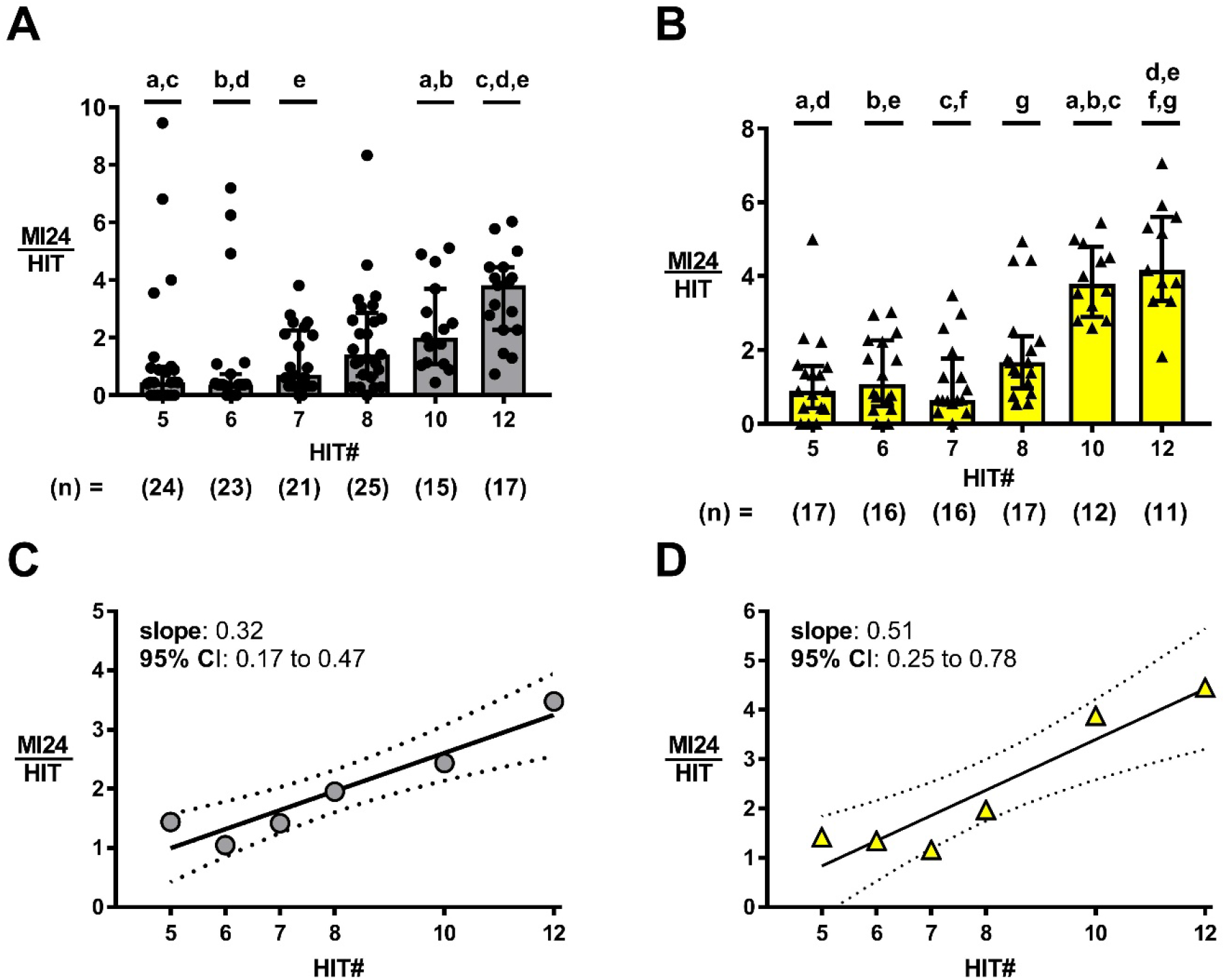
rmTBI causes a synergistic effect on mortality. **(A,B)**Mortality per injury is greater for both *w^1118^* (A) and *y^1^w^1^* (B) flies subjected to 10 or 12 HITs vs lesser injury numbers. Data plotted are median with IQR. Indicated ‘n’ values and symbols reflect each vial of ≥ 40 flies. Datasets with shared letters are statistically different from each other (p < 0.05 by Kruskal-Wallis with Dunn’s Correction). **(C,D)**Overall MI24/HIT values across 5-12 HITs for both *w^1118^* (C) and *y^1^w^1^* (D) flies best fit a positive slope line. Symbols are the overall MI24/HIT for the respective condition. Solid lines indicate the best-fit line with 95% confidence bands plotted by dashed lines. n ≥ 710 flies per condition for *w^1118^;* n ≥ 537 flies per condition for *y^1^w^1^.*

Our data shows that mild TBI causes synergistic effects on mortality after a requisite number of injuries, possibly because cumulative cellular stress surpasses a critical threshold. We expect that many of the secondary injury pathways active in animals following our mild TBI (60°) methodology overlap with pathways previously reported (Katzenberger et al. 2015; Katzenberger et al. 2016; Anderson et al. 2018; Saikumar et al. 2020; Swanson, Rimkus, et al. 2020; Swanson, Trujillo, et al. 2020). However, the identity of any specific factor(s) which set or scale the sensitivity to rmTBI remain unknown. Identification of such factor(s) may allow us to develop diagnostic tools more sensitive to tracking rmTBI outcomes and lead to identification of genetic risk factors which contribute to disease following rmTBI.

## Methods

### Fly Husbandry and TBI Methodology

Stocks of *w^1118^* (BL 5905) and *y^1^w^1^* (BL 1495) were obtained from the Bloomington Drosophila Stock Center (Bloomington, Indiana, USA). Flies were maintained in a humidified 25°C incubator, on a 12H:12H light:dark cycle, and on a standard glucose-cornmeal-yeast food (Putnam et al. 2019).

Methods for TBI were based on those reported previously (Katzenberger et al. 2013; Putnam et al. 2019). Briefly, flies were collected using CO2 anesthesia (35 – 60 flies per vial), and subjected to TBI by 5 days after eclosion. TBI was administered using a modified high-impact trauma (HIT) device with a stopping point which limited the spring deflection to 60°. The vial was released and allowed to collide with a foam pad covered by a 1/16” rubber pad. Injuries were spaced 15 seconds apart and repeated until a total of 5-12 injuries were administered. Animals were hand-transferred to food vials and returned to the 25°C incubator until they were assessed for mortality 24 hours later. The mortality index at 24 hours (MI24) was calculated as: MI24 = number of flies dead at 24 hours/total number of flies * 100. MI24/HIT values were calculated by dividing the MI24 value by the administered HIT number.

### Statistics

MI24/HIT values for comparisons of mean ranks across 5-12 HITs were calculated for each vial of at least 40 flies. Conditions were compared by Kruskal-Wallis with multiple comparisons of mean ranks and Dunn’s correction at the level of α= 0.05 (GraphPad Prism 7).

The pattern of mortality was tested via line-fit of MI24/HIT data across HIT number. A single MI24 value obtained from full dead:total count data was divided by the HIT number to calculate the MI24/HIT value for each condition. MI24/HIT data points were then plotted across HIT number using the linear fit mode in the nonlinear regression analysis toolkit, fitted using the least squares fit mode, compared to a hypothetical slope of zero via the extra sum-of-squares F test at a level of α= 0.05, and plotted with the best-fit slope and the asymmetrically determined 95% confidence interval (CI) (GraphPad Prism 7).

## Funding

UWGB Start-Up Funds

UWGB Research Scholar

